# Identification of cell-type-specific marker genes from co-expression patterns in tissue samples

**DOI:** 10.1101/2020.11.07.373043

**Authors:** Yixuan Qiu, Jiebiao Wang, Jing Lei, Kathryn Roeder

## Abstract

**Motivation:** Marker genes, defined as genes that are expressed primarily in a single cell type, can be identified from the single cell transcriptome; however, such data are not always available for the many uses of marker genes, such as deconvolution of bulk tissue. Marker genes for a cell type, however, are highly correlated in bulk data, because their expression levels depend primarily on the proportion of that cell type in the samples. Therefore, when many tissue samples are analyzed, it is possible to identify these marker genes from the correlation pattern.

**Results:** To capitalize on this pattern, we develop a new algorithm to detect marker genes by combining published information about likely marker genes with bulk transcriptome data in the form of a semi-supervised algorithm. The algorithm then exploits the correlation structure of the bulk data to refine the published marker genes by adding or removing genes from the list.

**Availability and implementation:** We implement this method as an R package markerpen, hosted on https://github.com/yixuan/markerpen.

## 1 Introduction

Cell-type-specific (CTS) genes, also known as marker genes, are genes that are highly expressed in one cell type, but lowly expressed in other types. These genes, which define cellular identity, are key to the analysis of RNA transcriptional data. Knowledge of marker genes gives insights into the core set of genes whose expression is shared among all cells of a given type, and will fill critical gaps in our understanding of cell biology and possibly the cellular origins of pathologies (Kelley et al., 2018). Marker genes are used to annotate cell clusters (Kiselev et al., 2017), to study cellular composition of bulk tissues (Oldham et al., 2008; Xu et al., 2013; Kelley et al., 2018; Luecken and Theis, 2019), to estimate cell type fraction via deconvolution (Gaujoux and Seoighe, 2012; Zhong et al., 2013; Abbas et al., 2009; Newman et al., 2015; Avila Cobos et al., 2018), and to estimate CTS expression directly from bulk tissue (Wang et al., 2020a,b).

Because marker genes are defined by their strong differential expression among cell types, a common approach to identifying them is to conduct statistical tests on CTS transcriptome data, typically single-cell RNA sequencing (RNA-seq). Genes that have significant expression differences between one specific cell type and all others are regarded as marker genes for this type (Kiselev et al., 2017). Despite the obvious appeal of this direct approach, the availability of CTS transcriptome data is a great challenge for many studies. The cost for single-cell sequencing is generally high, and in some cases, viable cells are hard to obtain for tissues like human brain. Even if public data sets are available, they might not correspond well with the data in hand, being collected at a different developmental period or a different functional portion of the organ. Furthermore, there is a trade-off between sequencing depth and the number of cells that can be analyzed, and for this reason the resulting single-cell transcriptome is quite noisy. An alternative way to obtain reference transcriptome data is to use single-cell RNA-seq data from another species (Zeisel et al., 2015); however, the quality of the obtained marker genes based on data from a different species is questionable. To this end, there is a need for a reliable statistical technique for detecting marker genes that does not require well matched single-cell RNA-seq data.

The objective of this inquiry is to develop a method for identifying a set of marker genes that describe the expression of the cells that constitute a tissue sample directly from the bulk transcriptome. We will take advantage of the conjecture that marker genes identifying a common cell type are highly correlated in samples of bulk transcriptome data, because their expression levels depend primarily on the proportion of that cell class in each sample (Oldham et al., 2008; Kelley et al., 2018). Motivated by this insight, we develop a new algorithm called MarkerPen, short for **marker** gene detection via **pen**alized principal component analysis, to detect marker genes by combining prior marker information with bulk transcriptome data. MarkerPen is a semi-supervised algorithm that requires two pieces of information: a list of potential marker genes, typically obtained from the literature, past experience, or available single-cell RNA-seq data; and a bulk RNA-seq data set, viewed as a mixture of pure cells. The algorithm then exploits the bulk data to refine the published marker genes by adding and removing genes from the list.

In summary, MarkerPen is motivated by the following two key findings: (1) marker genes are statistically highly correlated under mild and sensible assumptions; (2) highly correlated genes can be detected by estimating the leading eigenvectors of the correlation matrix. We formulate the MarkerPen algorithm as a modified sparse principal component analysis (sparse PCA, Jolliffe et al., 2003; Zou et al., 2006; Zou and Xue, 2018), which simultaneously selects highly correlated genes and encodes prior information about markers into the model. Our simulation study and multiple data analyses of human brain transcriptomes demonstrate the superior performance of the proposed method.

## 2 Materials and methods

### 2.1 Related work

The MarkerPen algorithm follows the path of two pioneering publications, Xu et al. (2013) and Kelley et al. (2018), who noted that marker genes tend to be highly correlated in bulk tissue. MarkerPen solves the marker detection problem by making better use of bulk RNA-seq data. The motivation for these methods is straightforward: many tissues and subjects have been assessed for bulk tissue expression; the data tend to be of better quality; and collecting bulk data is less costly. Although bulk data alone do not provide CTS transcriptome information, they can be combined with prior knowledge of marker genes to improve the quality of published markers. For example, Xu et al. (2013) first obtained CTS genes in mouse brain as potential markers for human brain, and then performed co-expression network analysis on human brain bulk data to select highly correlated genes of each type as the refined marker genes. This method has shown good empirical results, but has the drawback that genes can only be removed from the candidate list, but not added from the complementary set. More recently, Kelley et al. (2018) applied a similar approach to the human brain transcriptome. They first built an unsupervised co-expression network for all genes, and then identified gene clusters that were maximally enriched with published markers. Each gene was then assigned a fidelity score for each cell type, as an indicator for the strength of association between the gene and the cell type. These scores, however, were based on the aggregation of multiple data sets, and hence the selected marker sets may be suboptimal for a specific study.

Both methods described above assume that marker genes tend to be highly correlated, which is an intuitive assumption supported empirically in numerous species (Oldham et al., 2008; Fertuzinhos et al., 2014; Ponomarev et al., 2010; Hilliard et al., 2012; Bakken et al., 2016; Hawrylycz et al., 2015), but lacks rigorous statistical justification. To resolve this shortcoming, in the supplementary material (Section S.1) we explicitly study the statistical properties of marker genes, and show that under weak assumptions the marker genes for the same cell type are highly correlated in the bulk data. Given this fact, we are then able to utilize the correlation structure to detect marker genes via the MarkerPen algorithm.

### 2.2 The MarkerPen algorithm

Because high mutual correlation is a necessary condition for marker genes, the first step of marker gene selection is to find a subset from the whole genome such that genes in this set are highly correlated with each other. If the true correlation matrix Σ is available, then such a goal can be achieved by computing PCA on Σ, as the eigenvectors of Σ, also known as factor loadings, indicate the contribution of each gene to form a gene group. In the case of a marker gene group, the eigenvector contains a few strong signals and a large number of small values, where the large coefficients correspond to highly correlated genes (Section S.2, Figure S1).

However, in practice, only the sample correlation matrix *S* is given, and *S* can be of very high dimension. Theoretical results show that conventional PCA is likely to fail in high dimensions (Johnstone and Lu, 2009; Jung and Marron, 2009), so in this case the sparse PCA method is preferred, which directly estimates a sparse eigenvector, meaning that most entries in this vector are zeros. Sparse PCA has many different variants, and in this article we consider the Fantope projection and selection algorithm (FPS, Vu et al., 2013), because it solves a convex optimization problem that has a global convergence guarantee. Let Γ_*p×d*_ denote the eigenvectors of Σ associated with the largest *d* eigenvalues, where *p* is the number of genes, and then FPS estimates the top-*d* projection matrix Π_*p×p*_ = ΓΓ^T^ by solving

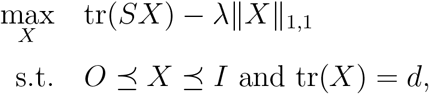

where tr(*A*) is the trace of a matrix *A*, ∥*X*∥_1,1_ = ∑_*i,j*_ |*X_ij_*| is the sum of absolute values of the elements in ***X***, λ is a tuning parameter that controls the sparsity of eigenvectors, and 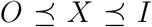 means all eigenvalues of *X* are between 0 and 1. Once we get an estimate 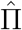 for the projection matrix Π, we can recover the eigenvectors Γ by computing the eigen decomposition of 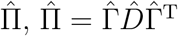.

In practice, there is abundant prior information about the marker gene list in the literature, which provides useful knowledge about the relationship between cell types and genes; however, such information is not exploited by FPS, resulting in low utilization of the available information. To fix this issue, the proposed MarkerPen algorithm modifies the original FPS such that prior information about markers can be combined with the collected bulk data. For simplicity, we first consider the detection of marker genes for one cell type. Let *G* be the indices of published marker genes for a cell type *C*, and then we solve

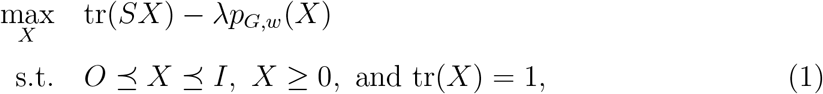

to estimate the projection matrix Π = *γγ*^T^, where *γ* is the leading eigenvector, 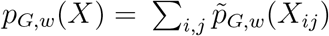 is a penalty function defined as

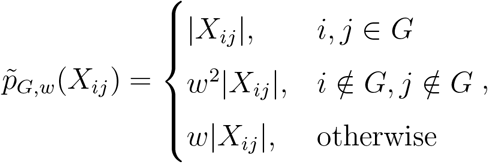

and *X* ≥ 0 means all elements of *X* are nonnegative. The added constraint *X* ≥ 0 is based on the fact that marker genes are positively correlated, so both the eigenvector *γ* and the projection matrix Π have nonnegative entries. The extra tuning parameter *w* ≥ 1 is used to put larger sparsity penalty on genes that are not in the prior list *G*, so that genes outside *G* are less likely to be selected as marker genes, unless they show large signals. The optimization problem (1) can be solved via the proximal-proximal-gradient method (Ryu and Yin, 2017), with details in the supplementary material (Section S.3).

After we obtain the estimate for the leading eigenvector *γ*, we select genes that have coefficients greater than some threshold *ε* > 0, and treat them as marker genes for cell type *C*. For multiple cell types *C*_1_, *C*_2_,…, we repeatedly apply the algorithm above to compute different marker gene groups sequentially.

### 2.3 Data sources

In the next section we validate the performance of MarkerPen using a broad range of bulk and single-cell RNA-seq data, and here we provide some basic information of each data set. Below are the bulk tissue data used in this article:

1. **MSBB** The Mount Sinai/JJ Peters VA Medical Center Brain Bank cohort (Wang et al., 2018) contains RNA-seq data from human temporal cortex, with 425 control samples and 425 samples from patients with Alzheimer’s disease (AD, Braak score ≥ 4). Only the control samples are used.
2. **ROSMAP** The Religious Orders Study and the Rush Memory and Aging Project (Mostafavi et al., 2018; De Jager et al., 2018) collects RNA-seq data from the human dorsolateral prefrontal cortex (DLPFC), with 288 control samples and 348 AD samples. Only the control samples are used.
3. **MayoRNAseq** The Mayo Clinic RNA-seq data set (Allen et al., 2016; Allen et al., 2018) contains human temporal cortex RNA-seq data with 28 control samples and 82 AD samples. Only the control samples are used.
4. **BrainVar** The BrainVar data set (Werling et al., 2020) consists of 176 samples from the human DLPFC across development, from 6 post-conception weeks to young adulthood. To be comparable with other data sets we exclude pre-natal brains and focus on subjects that are at least 6 months old (epoch 3), finally with a sample size of 45.
5. **CMC** The human brain RNA-seq data collected by the CommonMind Consortium (Fromer et al., 2016) contain 258 adult schizophrenia subjects and 279 adult control subjects, and only the control samples are used. As the original data set spans a broad range of ages, we further split the control group into two subsets, resulting in groups with ages less than or equal to 70 (sample size 164) and greater than 70 (sample size 115).

We also use single-cell and single-nucleus RNA-seq data sets:

1. Mathys et al. (2019) provides single-nucleus transcriptomes from DLPFC of 48 subjects with varying degrees of AD pathology. Only the data from 17 control subjects are used.
2. Darmanis et al. (2015) obtains single-cell RNA-seq data of human cortical tissues from eight adults and four embryonic samples. Only the adult data are used.
3. Li et al. (2018) collects single-nucleus RNA-seq data from DLPFC of three adult brains.
4. Zeisel et al. (2015) provides mouse cerebral cortex single-cell RNA-seq data.

## 3 Results

### 3.1 Quality of selected markers

In this section we demonstrate the quality of marker genes selected by MarkerPen from three different angles.

First, as explained in Section 2.1, we expect to see that marker genes for the same cell type are highly correlated in the bulk data. Therefore, the quality of selected marker genes can be visually examined by the correlation matrix. We study human brain bulk tissue RNA-seq data, and use the MSBB data set for illustration. To apply the MarkerPen algorithm, the prior marker gene list is obtained from existing literature, including 184 marker genes for astrocytes, 130 genes for oligodendrocytes, 319 genes for neurons (all three from Cahoy et al., 2008), 100 genes for microglia (Hickman et al., 2013), and 237 genes for endothelial cells (Butler et al., 2016). Figure 1A shows the sample correlation matrix of the published marker genes in the MSBB bulk data. It can be seen that the correlation matrix roughly forms five blocks, but the boundary between the blocks is not very clear as much noise exists.

**Figure 1:**
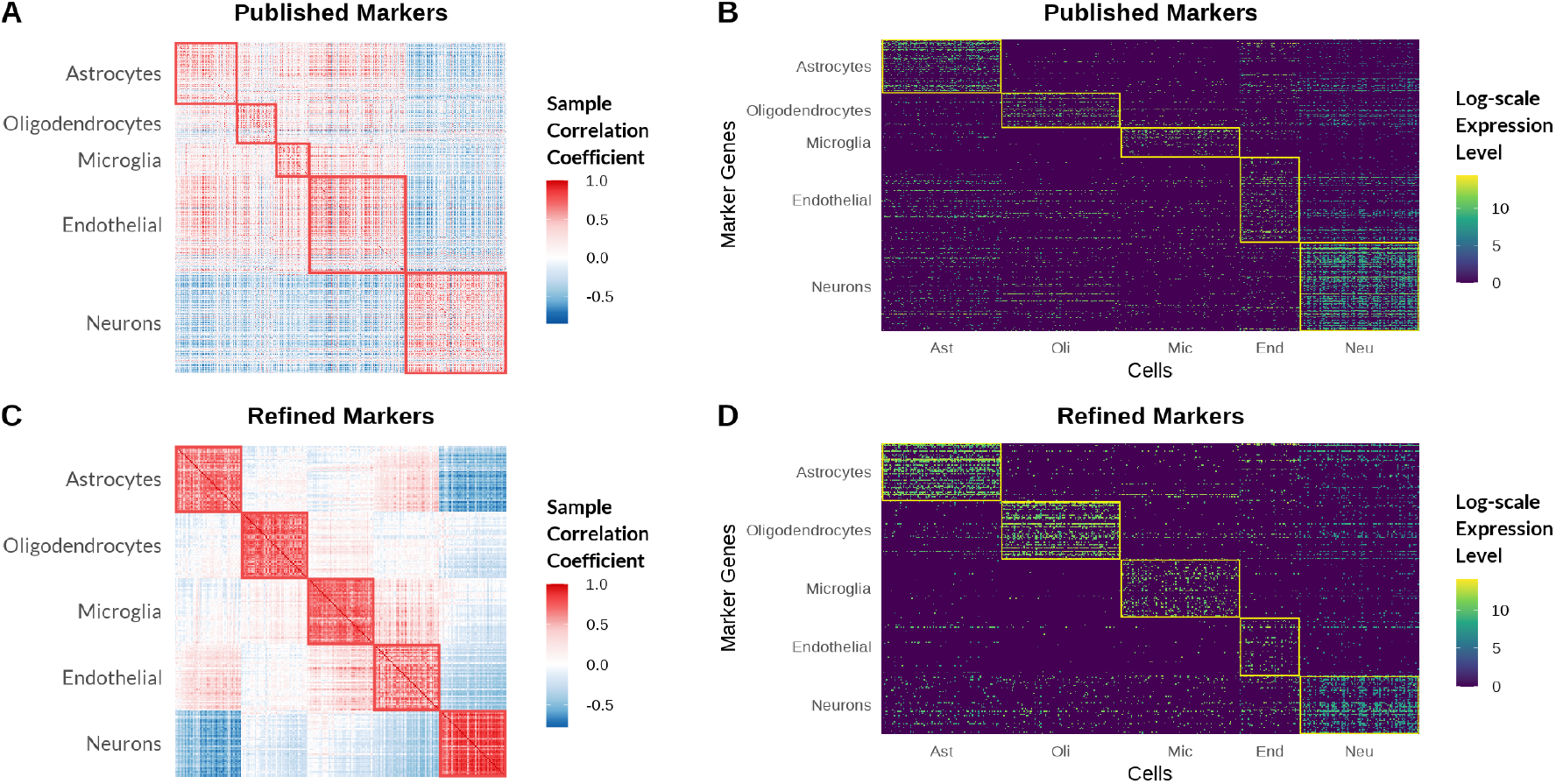
(A) Sample correlation matrix of published marker genes in the MSBB bulk data. (B) Gene expression of single-cell reference data from Mathys et al. (2019) on published marker genes. (C) Sample correlation matrix of refined marker genes output by MarkerPen. (D) Gene expression of single-cell reference data on refined marker genes.

Then we apply the MarkerPen algorithm to refine the given marker gene list. For each cell type, we restrict the search range to the union of the published marker genes and the top 500 genes that have the highest fidelity scores given by Kelley et al. (2018). Figure 1C demonstrates the sample correlation matrix of the refined genes, in which 50 genes are selected for each cell type. It is clear that after the refinement, genes in the same block have much stronger mutual correlation, whereas genes in different blocks are only weakly correlated. In other words, genes refined by MarkerPen have a correlation structure that better fits the property of marker genes.

Second, by definition, marker genes should be largely expressed in one cell type but weakly expressed in others. Therefore, it is helpful to examine the expression level of selected marker genes in purified single-cell data. We use the single-nucleus transcriptome data from Mathys et al. (2019) to demonstrate this idea. For each cell type, we randomly select 100 samples (50 for endothelial due to the limited number in the data set), and plot the logarithm-scale expression matrix on published and refined marker genes in Figure 1B and D, respectively. In Figure 1B, we can observe that many genes in the published list behave like noise, as they show very low expression level in virtually all cell types. In contrast, this defect has been greatly reduced in Figure 1D, where most noise genes have been removed by MarkerPen. This finding further justifies the MarkerPen selection algorithm.

Finally, considering that the transcriptome data from Mathys et al. (2019) and the MSBB bulk data may not fully match, it is more appropriate to study the purified cells from the same subjects as in the bulk data. However in practice, this is not always possible. Instead, we can use the bMIND algorithm (Wang et al., 2020b) to estimate CTS gene expression for each subject in the bulk data. The output of bMIND can be viewed as the average of denoised single-cell data for the subjects in the bulk data. We plot the estimated CTS gene expression matrix on three types of markers: the published marker genes, the markers selected by MarkerPen, and the bMIND markers that are directly selected from the estimated CTS gene expression. The bMIND markers are treated as the ground truth. However, because marker genes form a highly correlated set, there is not a unique set of optimal genes to serve this purpose. In our evaluation we look to see if the set of selected markers matches the good properties exhibited in the bMIND set. The first row of Figure 2 demonstrates the results for the MSBB data set, from which we can find that published markers contain a lot of noise, whereas the MarkerPen output is very similar to that of bMIND. Also included in Figure 2 are the results for two additional bulk data sets: the ROSMAP and MayoRNAseq data. They both give similar results that validate the quality of MarkerPen genes.

**Figure 2:**
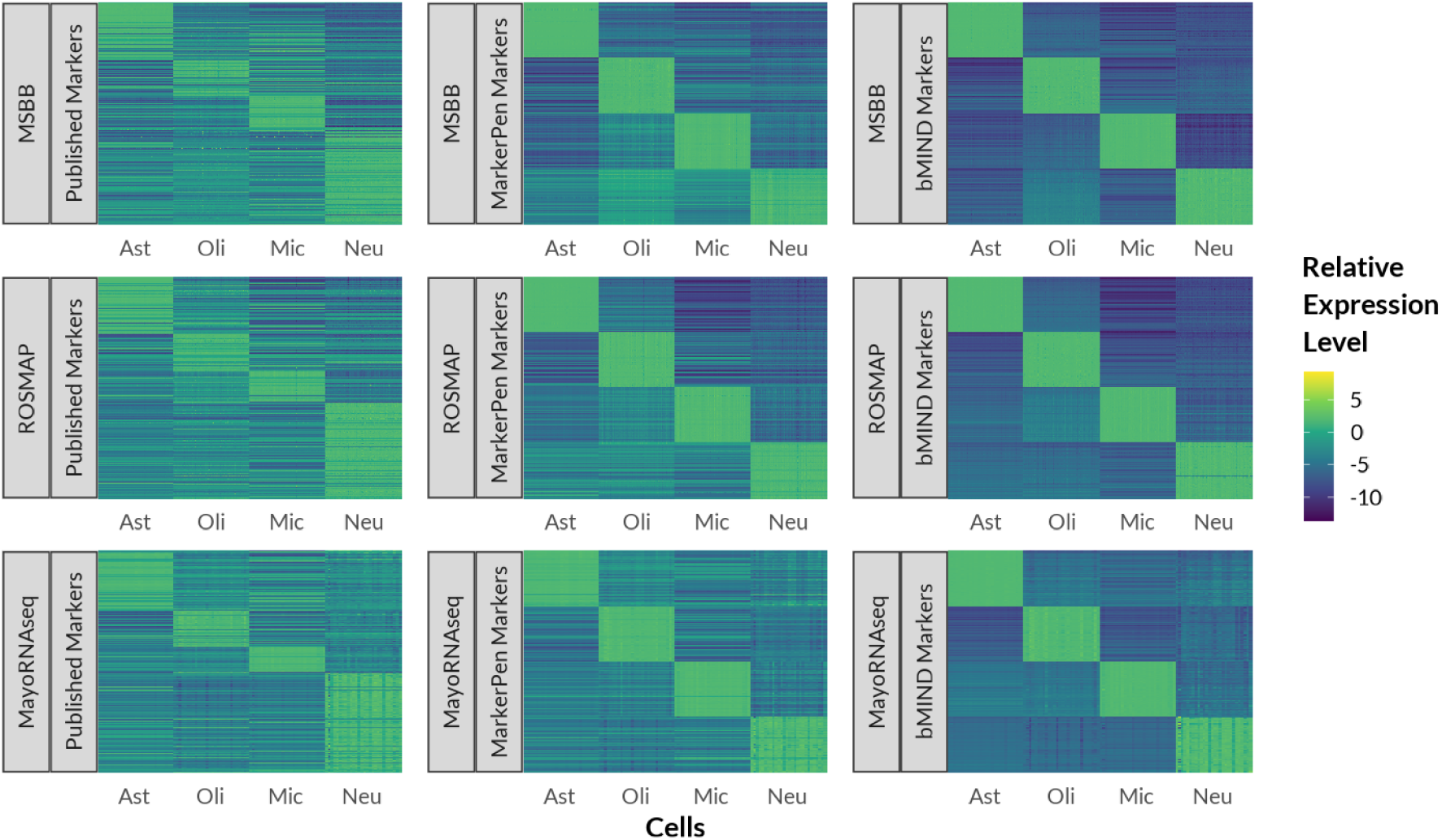
CTS gene expression of the MSBB, ROSMAP, and MayoRNAseq data sets on three types of marker genes: the published markers, the ones selected by MarkerPen, and the bMIND markers that can be treated as the truth. Ast=astrocytes, Oli=oligodendrocytes, Mic=microglia, Neu=neurons.

### 3.2 Performance in downstream analysis

As marker genes are essential tools for many downstream analyses such as cell type fraction deconvolution, in this section we use simulation experiments to evaluate the performance of our algorithm in such tasks. Cell type fraction deconvolution is a problem commonly seen in bulk RNA-seq data analysis. Because the deconvolution result depends on the selection of marker genes, the quality of the selected markers can be measured by the estimation error of cell type fractions. We design a simulation experiment to compare MarkerPen with two supervised marker gene selection algorithms, with experiment setting described in the supplementary material (Section S.4, Figure S2, S3).

In practice, deconvolution can be conducted with or without single-cell reference samples, and the quality of reference samples may also vary. To reflect these different scenarios, we design three models for simulating the observed data:

1. **Matched reference case** Reference samples and the bulk data are simulated from the same signature matrix.
2. **Noisy reference case** The bulk data use a perturbed version of the signature matrix: some percentage of the genes, ranging from 5% to 30%, are set to noise. This indicates that some genes may be markers in the reference data, but they play no role in the bulk data.
3. **No reference case** No reference samples are simulated.

For model 1 and model 2, both the bulk data and the reference samples are available, and we use a supervised method, **dtangle** (Hunt et al., 2018), to accomplish the deconvolution. For model 3, only the bulk data and the marker gene list are available, so we apply a semi-supervised algorithm for deconvolution, the digital sorting algorithm (**DSA**, Zhong et al., 2013). The choice of deconvolution algorithms is beyond the scope of this article, as the main purpose of this section is to evaluate the effect of marker gene selection for a fixed deconvolution method. In practice any deconvolution algorithm that needs marker genes can be used in place of the methods investigated here.

In our experiments, we use the mouse brain single-cell RNA-seq data from Zeisel et al. (2015) to simulate the true signature matrix. We select seven major cell types (astrocytes, oligodendrocytes, microglia, endothelial, interneurons, S1 pyramidal neurons, and CA1 pyramidal neurons) from the whole single-cell data, and restrict to 2452 genes that are known to be associated with the cell types (Table S1 of Zeisel et al., 2015). Following the steps in Section S.4, we simulate the fraction matrix, reference samples, and the bulk data according to a stochastic model. From the signature matrix, we randomly select 50 genes from each cell type block, and treat them as known marker genes. Of course, due to the possible perturbation of the signature matrix, some of the claimed marker genes will be noise in the bulk data, and hence provide little information about the cell type. This treatment is used to mimic the quality of marker genes in reality.

We repeat the procedure above 30 times, so that in every simulation run, the generated data are different but follow the same stochastic model. We compute the deconvolution estimation errors in each simulation run, and summarize their distribution density curves in Figure 3.

**Figure 3:**
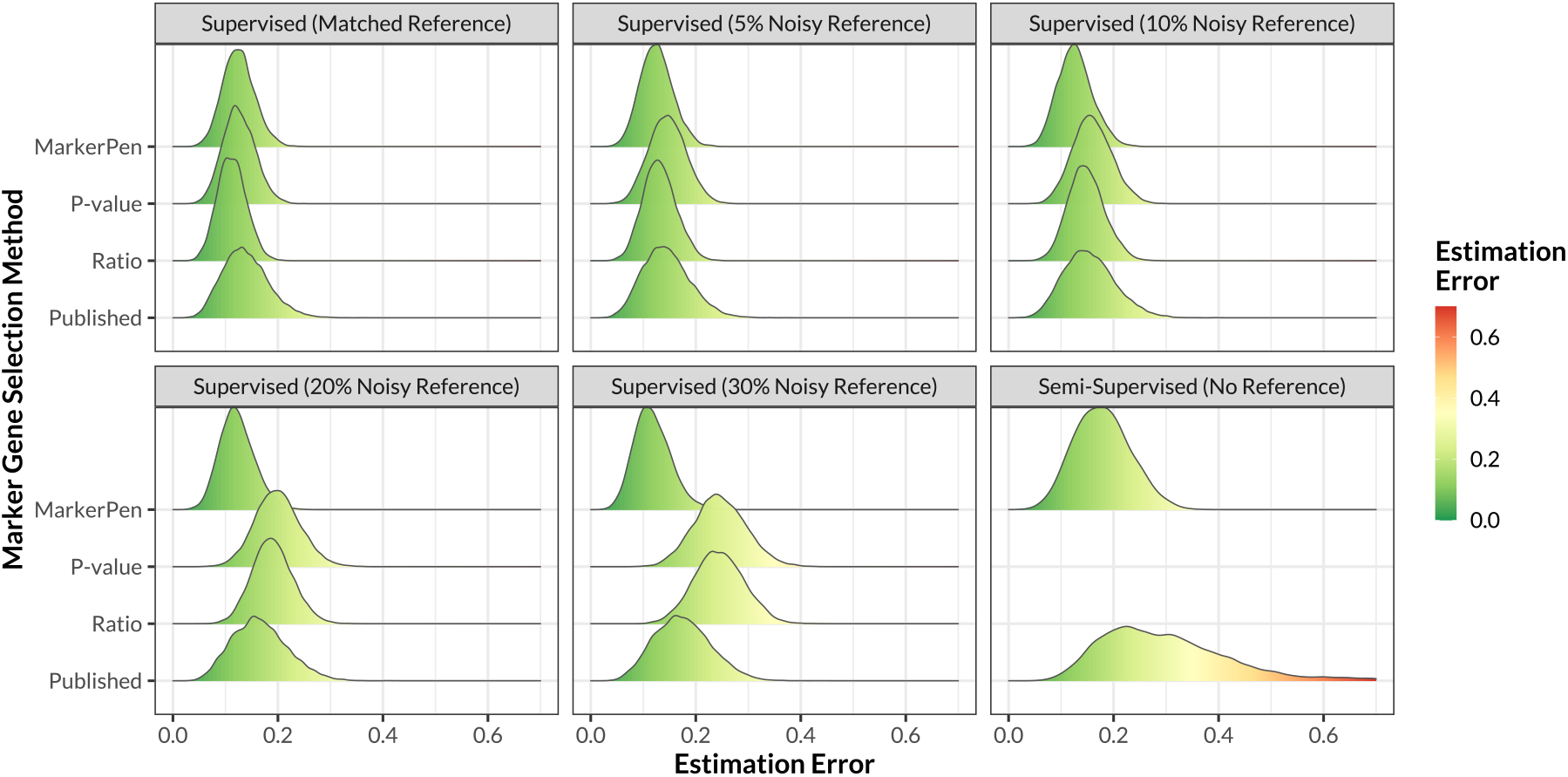
Impact of marker gene selection algorithm on deconvolution estimation error, displayed as distribution density curves. The vertical axis stands for different marker gene selection methods. MarkerPen: the proposed method. P-value and Ratio: selection methods based on reference samples, implemented in the **dtangle** R package. Published: using all published marker genes without selection.

In Figure 3, each panel represents one model for the reference sample. It is clear that when the reference sample and bulk data are matched, all marker gene selection methods behave equally well, compared with the last row that stands for no selection. However, when the noise level increases, selection methods purely based on the reference sample become much worse, whereas the proposed MarkerPen is quite robust and accurate. When no reference sample is available, reference-based selection methods do not apply, but MarkerPen still shows improvement via semi-supervised marker gene selection. These findings highlight the power of MarkerPen in refining published marker genes.

### 3.3 Robustness

In Section 3.2 we have studied the accuracy of MarkerPen in downstream deconvolution tasks. Then a natural question is how robust MarkerPen is across different data sets. To answer this question, we experiment on the combination of four bulk data sets and three single-cell and single-nucleus reference data sets, and study the variation of their deconvolution results. Descriptions of these data sets are given in Section 2.3.

For each pair of data sets, we estimate the cell type fractions for each observation, using three marker gene selection methods: the proposed MarkerPen, the supervised method based on single-cell or single-nucleus reference data, and a fixed set of marker genes given by the **BRETIGEA** R package (McKenzie et al., 2018). Figure 4A shows the estimated fractions averaged over all observations in the data set. It is easy to see that the supervised algorithm and **BRETIGEA** generate significantly different results under three reference data sets, whereas MarkerPen is much more consistent and robust. We then compute a metric (Section S.5) to measure the variation of estimated fractions across different reference data sets, and show the values in Figure 4B. The first four panels give the comparison in each bulk data set, and the last panel shows the result over all data sets. In all settings MarkerPen is much more robust to the choice of single-cell reference data compared with others.

**Figure 4:**
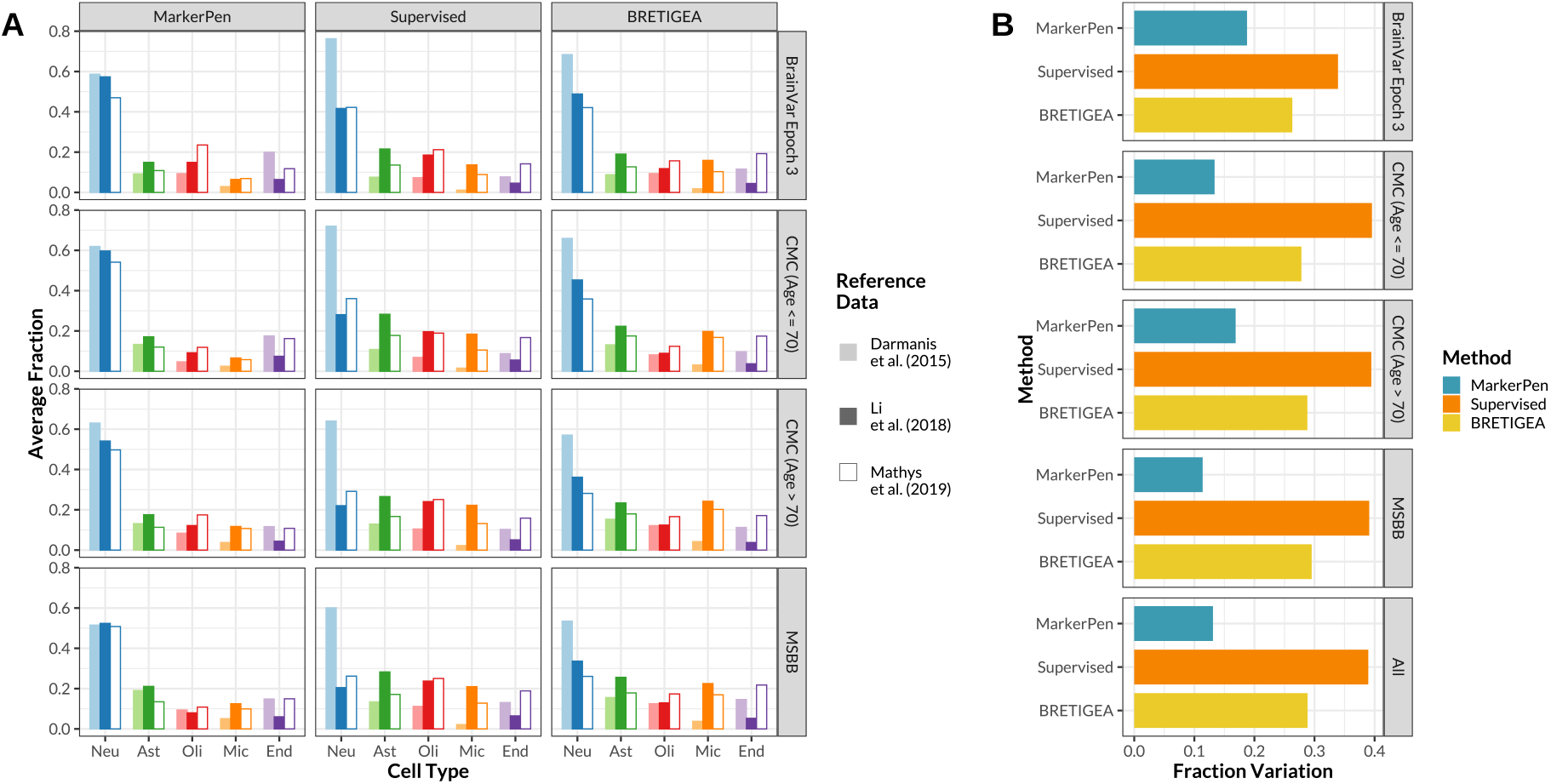
(A) Estimated average cell type fraction on different bulk data sets using three single-cell reference data sets and three marker gene selection methods. Deconvolution is conducted using the **dtangle** package. Neu=neurons, Ast=astrocytes, Oli=oligodendrocytes, Mic=microglia, End=endothelial. (B) Comparison of the fraction variation metric for three marker gene selection methods under various bulk data sets. This metric is used to quantify the variation of fraction estimates across different single-cell and single-nucleus reference data sets.

## 4 Conclusion and discussion

We have presented the MarkerPen algorithm for identifying cell-type-specific marker genes from bulk tissue data. Unlike most marker gene detection methods that heavily rely on single-cell reference samples, MarkerPen is a semi-supervised method that only requires the bulk data and a prior marker gene list. This feature makes the algorithm especially useful when tissue level data are not well matched with available single-cell data. More importantly, using well selected marker genes corrects the bias and error of downstream analyses of bulk tissue samples. Furthermore, MarkerPen interfaces nicely with other marker gene selection algorithms. For example, supervised methods applied to single-cell RNA-seq data can provide the prior gene list for MarkerPen.

A promising application of MarkerPen is to study the evolution of marker genes over developmental stages. Preliminary studies of the CMC data reveal that some marker genes identified from younger subjects are less correlated in older brains. The BrainVar data, which include brains sampled over all developmental stages, would provide an ideal data set to further investigate how marker genes change over time; however, it will be more challenging to compare marker genes of mature brains with those of fetal brains. We leave this topic for future explorations.

The use of single-cell RNA-seq has increased. However, there are drawbacks to single-cell data, including its noisy nature and the limited number of subjects from whom cell are harvested for study. By contrast, bulk transcriptome data are less noisy, and they can readily be sampled from many subjects at a reasonable cost. With larger sample sizes, bulk tissue samples can be much more informative for downstream analyses, such as eQTL mapping. With the help of good marker genes, many deconvolution methods can provide accurate estimates of cell type fractions (Zhong et al., 2013; Gaujoux and Seoighe, 2013; Newman et al., 2015; Hunt et al., 2018; Newman et al., 2019). Furthermore, cell type fractions are input of methods such as MIND (Wang et al., 2020a) and bMIND (Wang et al., 2020b) to estimate CTS expression profiles from bulk tissue samples, permitting cell-type analysis for features such as eQTLs. The performance of these algorithms is highly dependent on the selection of good marker genes, hence MarkerPen can play a critical role in the analysis of CTS expression.

There are two limitations to the current version of MarkerPen. First, although MarkerPen is based on the eigen decomposition of correlation matrices, its computational complexity is greater than ordinary principal component analysis. In practice, one might need to limit the search range of genes to a few thousand. Despite this restriction, the algorithm has been implemented in the **markerpen** R package with core part written in efficient C++ code. Another challenge for MarkerPen is to detect cell types that are similar, such as neuron subtypes. These subtypes do not induce a strict block structure in the correlation matrix, making it harder to identify subtype-level marker genes.

MarkerPen can be extended in several directions. For instance, the current algorithm that selects marker genes performs the calculation on one cell type at a time. It may achieve better performance, however, by jointly selecting mutually exclusive marker genes for multiple cell types. Another promising direction would be to extend MarkerPen to analyzing unannotated single-cell RNA-seq data. It might be useful in selecting marker genes for clustering unlabeled cells.

## Acknowledgments

We are indebted to Bernie Devlin for suggesting this topic of inquiry and Lu Xie for preliminary investigations of the idea several years ago. We benefited from comments on data analysis from Gabriel Hoffman, Michael Breen and Panos Roussos.

Data were generated as part of the CommonMind Consortium supported by funding from Takeda Pharmaceuticals Company Limited, F. Hoffman-La Roche Ltd and NIH grants R01MH085542, R01MH093725, P50MH066392, P50MH080405, R01MH097276, RO1-MH-075916, P50M096891, P50MH084053S1, R37MH057881, AG02219, AG05138, MH06692, R01MH110921, R01MH109677, R01MH109897, U01MH103392, and contract HHSN271201300031C through IRP NIMH. Brain tissue for the study was obtained from the following brain bank collections: the Mount Sinai NIH Brain and Tissue Repository, the University of Pennsylvania Alzheimer’s Disease Core Center, the University of Pittsburgh NeuroBioBank and Brain and Tissue Repositories, and the NIMH Human Brain Collection Core. CMC Leadership: Panos Roussos, Joseph Buxbaum, Andrew Chess, Schahram Akbarian, Vahram Haroutunian (Icahn School of Medicine at Mount Sinai), Bernie Devlin, David Lewis (University of Pittsburgh), Raquel Gur, Chang-Gyu Hahn (University of Pennsylvania), Enrico Domenici (University of Trento), Mette A. Peters, Solveig Sieberts (Sage Bionetworks), Thomas Lehner, Stefano Marenco, Barbara K. Lipska (NIMH).

The results published here are in part based on data obtained from the AD Knowledge Portal (https://adknowledgeportal.synapse.org). Study data were provided by the Rush Alzheimer’s Disease Center, Rush University Medical Center, Chicago. Data collection was supported through funding by NIA grants P30AG10161 (ROS), R01AG15819 (ROSMAP; genomics and RNAseq), R01AG17917 (MAP), R01AG30146, R01AG36042 (5hC methylation, ATACseq), RC2AG036547 (H3K9Ac), R01AG36836 (RNAseq), R01AG48015 (monocyte RNAseq) RF1AG57473 (single nucleus RNAseq), U01AG32984 (genomic and whole exome sequencing), U01AG46152 (ROSMAP AMP-AD, targeted proteomics), U01AG46161(TMT proteomics), U01AG61356 (whole genome sequencing, targeted proteomics, ROSMAP AMP-AD), the Illinois Department of Public Health (ROSMAP), and the Translational Genomics Research Institute (genomic). Additional phenotypic data can be requested at www.radc.rush.edu.

